# In-silico analysis of important mitochondrial microRNAs and their differential expression in mitochondria

**DOI:** 10.1101/2024.04.25.591201

**Authors:** Ashutosh Kumar Maurya, P Rabina, V.B. Sameer Kumar

## Abstract

Mitochondria, often called as the power house of cell, serves important role in cellular physiology and survivability. It plays crucial role in the normal functioning of the cell. Dysfunctional mitochondria have been found to be associated with various pathological conditions including cancer. The modulation of mitochondrial machinery could be due to the suppression of the expression pattern of important mitochondrial genes and microRNAs could be considered as the key player in reprograming of the mitochondrial metabolism. Apart from the microRNAs coded by mitochondrial genome, nuclear coded microRNAs gets localized to the mitochondria and they influence the mitochondrial machinery by targeting the important mitochondrial genes. This group of microRNAs are called mitochondrial miRNAs or MitomiRs. In this study we selected 10 important candidate mitochondrial microRNAs and checked their abundance in the cancerous and non cancerous hepatic cell line (HepG2 and WRL68), followed by their differential expression in the mitochondria of the respective cell line. The results shown an inverse relation in the expression pattern of the candidate microRNAs with mitochondrial target genes, suggesting their direct targeting, as predicted by our in-silico data.

## 1. Introduction

Continuous energy supply to the cancer cells is a prerequisite for their indefinite growth and proliferation (Feitelson MA et.al, 2015). To accomplish this, cancer cells modify their mitochondrial metabolism (Shahruzaman SH et. al,2018). This alteration of mitochondria metabolism could result due to the suppression of expression of important mitochondrial genes involved in ATP synthesis (Dorji, J. et.al, 2020, Li J et.al, 2020). MicroRNA, a class of non-coding RNAs, could be considered as the key players in this metabolic switch (Zhang et.al, 2021), as they target the important mitochondrial genes involved in the energy production (Ao X et.al, 2023). They regulate the expression pattern of a variety of genes involved in normal cellular functioning, by a mechanism similar to RNA interference (Gulyaeva LF, 2016). MicroRNAs could alter the expression of a gene either at transcriptional or translational level (Fabien M et.al, 2010). Along with regulating the expression of the genes involved in regular functioning (Saliminejad K et.al, 2019), a variety of microRNAs have been reported to be associated with various disease conditions including cardiac disease, neurodegenerative disease, and cancer (Suzuki H, 2023, Ali Syeda et.al, 2020, Saliminejad K et.al, 2019). The microRNAs that promote carcinogenesis are called oncomiRs and they specifically target the tumor suppressor genes (Otmani K et.al,2022). Similarly, the miRNAs, which inhibit the expression of oncogenes, are called tumor suppressor microRNA (Sylvia K et.al, 2009). Various studies have shown the involvement of microRNAs in various types of cancers including breast cancer, colon cancer, hepatocellular carcinoma etc. (Reddy K.B. et. al, 2015).

In this study we tried to elucidate the variation in the pattern of microRNA levels in the mitochondria of cancerous and non cancerouse hepatic cells (HepG2 and WRL cells). As part of this study, we selected the 10 most important microRNAs on the basis of their nature i.e. oncogenic or tumor suppressive, involvement in various fatal cancers and most important criteria used was that, all of these candidate microRNAs were nuclear coded and their role as mitochondrial microRNAs in regulation of mitochondrial machinery was not yet reported. Following this, we performed detailed bioinformatic analysis of candidate microRNAs to predict their mitochondrial target genes using 5 different database (i.e. MicroRNA.org, Target Scan, miRDB, miRanda, miRBase), followed by the scoring analysis. We selected 5 highest scored mitochondrial genes ((ND6, ATP6, Cyto-B, Cox1, ND4L), involved in the mitochondrial metabolism to check for their differential expression in HepG2 and WRL cell lines. Parallely, we performed comparative analysis of the expression of candidate microRNAs in the clinical samples of the hepatic cancer at the initial and advance stage, to identify the important microRNAs playing role in hepatic cancer.

Next we checked the expression level of candidate miRNAs in cancerous and non cancerous hepatic cells, following which their enrichment in the mitochondria was checked. Further, we checked the expression level of the mitochondrial genes targeted by the candidate miRNAs to verify if the miRNA mediated regulation is a possibility.

## 2. Methodology

### Cell culture

Hela (cervical cancer cell line), WRL-68 (Human hepatic non cancerous cell line) and HepG2 cells (Hepatic carcinoma cell line) were cultured in DMEM supplemented with 10% FBS, antibiotic-antimycotic solution and L-Glutamine. The cells were maintained under standard culture conditions at 37°C with 5% CO2 and 95% humidity. For experiments, seeding density of 0.4 × 10^4^ cells (96 well), 0.6×10^6^ cells (30mm dish), 0.8×10^6^ cells (60mmdish), 2.2×10^6^ cells (100 mm dish) were used.

### Isolation of mitochondria

The mitochondria were isolated from the cells using hypotonic buffer, where the cells were allowed to swell in the buffer for 10 minutes and then break open the cells to release the mitochondria. The cell suspension was then centrifuged at 1300g to remove the cell debris, followed by centrifugation at 12000g to get the mitochondrial pellet. The mitochondrial pellet was suspended in the mitochondrial resuspension buffer.

### Sonication

The mitochondrial pellet was mixed with Lysis buffer and sonicated for 2 minutes at 70% amplitude with 15 sec ON and 30 sec OFF cycle on 4°C. The solution obtained, was centrifuged at 12000g for 10 minutes. The supernatant was collected and protein estimation was done followed by sample preparation for SDS PAGE.

### Protein estimation

Protein level of mitochondria was estimated by Bradford method (Bradford etal,1976). To achieve this, 10μl of sample and 90μl of bradford reagent (50 mg Coomasie Brilliant Blue-G250 in 25ml ethanol and 50ml of phosphoric acid made upto 100ml with water) was added in triplicates in 96 well plate and the absorbance was taken at 595nm by multimode plate reader. The concentration of protein was calculated from the standard plot to BSA with concentration range from 10μg-100μg.

### SDS-PAGE

Protein sample was prepared by mixing of 6x SDS loading dye and boiling it at 90°C for 10 minutes in water bath. The sample was immediately kept on ice and briefly centrifuged before loading on SDS-PAGE gel. The electrophoresis was carried out by using Bio-Rad electrophoresis unit. The protein samples were run through the stacking gel at 80V for 15 minutes and through the resolving gel at 100V at room temperature until the dye reached the end of the gel.

### Western blot analysis

The purity of the mitochondrial pellet was checked by western blot using mitochondria specific antibody (VDAC). Also, the mitochondrial pellet was checked for the nuclear and cytoplasmic contaminants using Histone H3 antibody for Nucleus and Hexokinase HK3 antibody for the cytoplasm.

### RNA Isolation

RNA was isolated from the mitochondrial pellet as well as from the total cell using trizole reagent. Following this, the concentration of the RNA was checked by using nano drop.

### Polyadenylation of RNA

Poly A tail was added to the RNA by Poly A Polymerase enzyme, using manufacturers protocol. This reaction set up was incubated at 37°C for 30 minutes followed by heat inactivation for 5 minutes at 65°C.

### cDNA synthesis

The polyadenylated RNA were used for the synthesis of miRNA specific cDNA by Kang method. Apart from this, total RNA was used to synthesize the cDNA for checking the expression of mitochondrial genes.

### Real Time PCR (qRT PCR)

Quantative real time PCR was performed to check the expression pattern of the candidate microRNA and other mitochondrial genes in mitochondria before and after over expression of candidate miRNAs. RNU6B gene was taken as internal reference gene for microRNA expression study and GAPDH was taken as internal control for mitochondrial genes.

#### Selection of candidate microRNAs

The microRNAs were selected on the basis of their nature i.e. oncogenic/tumor suppressive, their involvement in various common cancers and which have not been reported in alteration of ETC. Scientific literature data base, Pubmed was used for detailed survey of all the studies done till date involving the candidate microRNAs with our selection criteria and all together 215 publications were retrieved and reviewed. With literature survey, we found that, though the candidate microRNAs were reported to be involved in initiation and progression of various types of cancer, we could not find any study suggesting their role in modulating the mitochondrial metabolism.

#### In-silico analysis of clinical samples of HCC

47 clinical samples of HCC were analyzed for estimating the relative expression of candidate microRNAs by comparing the expression at initial stage with the advance stage of HCC. Gene Expression Omnibus (GEO) database (Accession ID-GSE123972), was used for this study.

### Target prediction and scoring

Target prediction analysis and scoring of candidate microRNA was done for mitochondrial genes, using five algorithms i.e. MirBase, miRanda, Target Scan, miRDB and MicroRNA.org. Five highest scored mitochondrial genes (ND6, ATP6, Cyto-B, Cox1, ND4L) were selected for target validation.

### Statistical analysis

All the data in the study were expressed as the mean with the standard error mean of at least three experiments, each done in triplicates. SPSS 11.0 software was used for analysis of statistical significance of difference by Duncan’s One way Analysis of Variance (ANOVA). A value of P<0.05 was considered significant.

## 3. Results

### Selection of 10 most important miRNAs

The microRNAs were selected by referring literature available till date, on the basis of their oncogenic nature, involvement in important cancers and which have not been reported to act as mitomiR or involved in regulating the Ox-Phos pathway of mitochondria.

**(Table 3.1)**

**Table 3.1:**
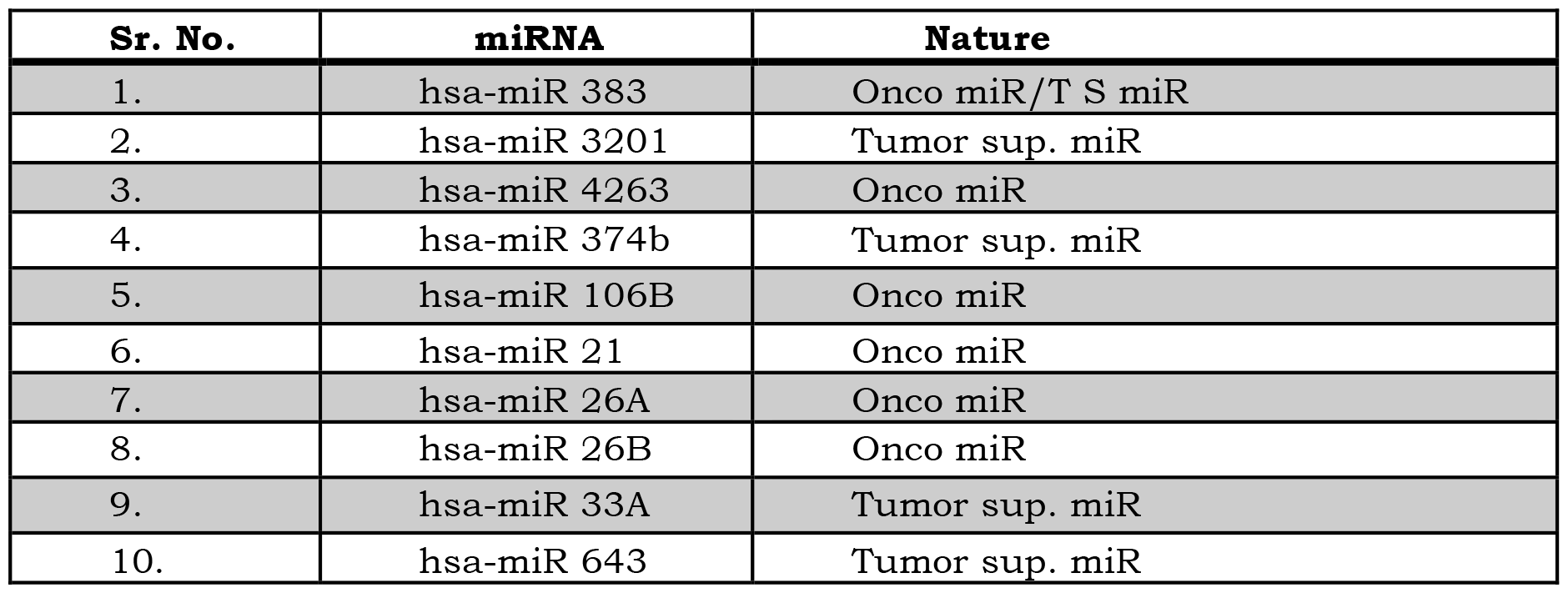
List of candidate microRNAs. The microRNAs were selected by referring literature available till date, on the basis of their nature, involvement in important cancers and which have not been reported to act as mitomiR or involved in regulating the Ox-Phos pathway of mitochondria.

| Sr. No. | miRNA | Nature |
| --- | --- | --- |
| 1. | hsa-miR 383 | Onco miR/T S miR |
| 2. | hsa-miR 3201 | Tumor sup. miR |
| 3. | hsa-miR 4263 | Onco miR |
| 4. | hsa-miR 374b | Tumor sup. miR |
| 5. | hsa-miR 106B | Onco miR |
| 6. | hsa-miR 21 | Onco miR |
| 7. | hsa-miR 26A | Onco miR |
| 8. | hsa-miR 26B | Onco miR |
| 9. | hsa-miR 33A | Tumor sup. miR |
| 10. | hsa-miR 643 | Tumor sup. miR |

### Target prediction and scoring analysis

The target prediction analysis was done using miRDB, MiRanda, Target scan, miRbase and microRNA.org.

**(Table 3.2)**

**Table 3.2:**
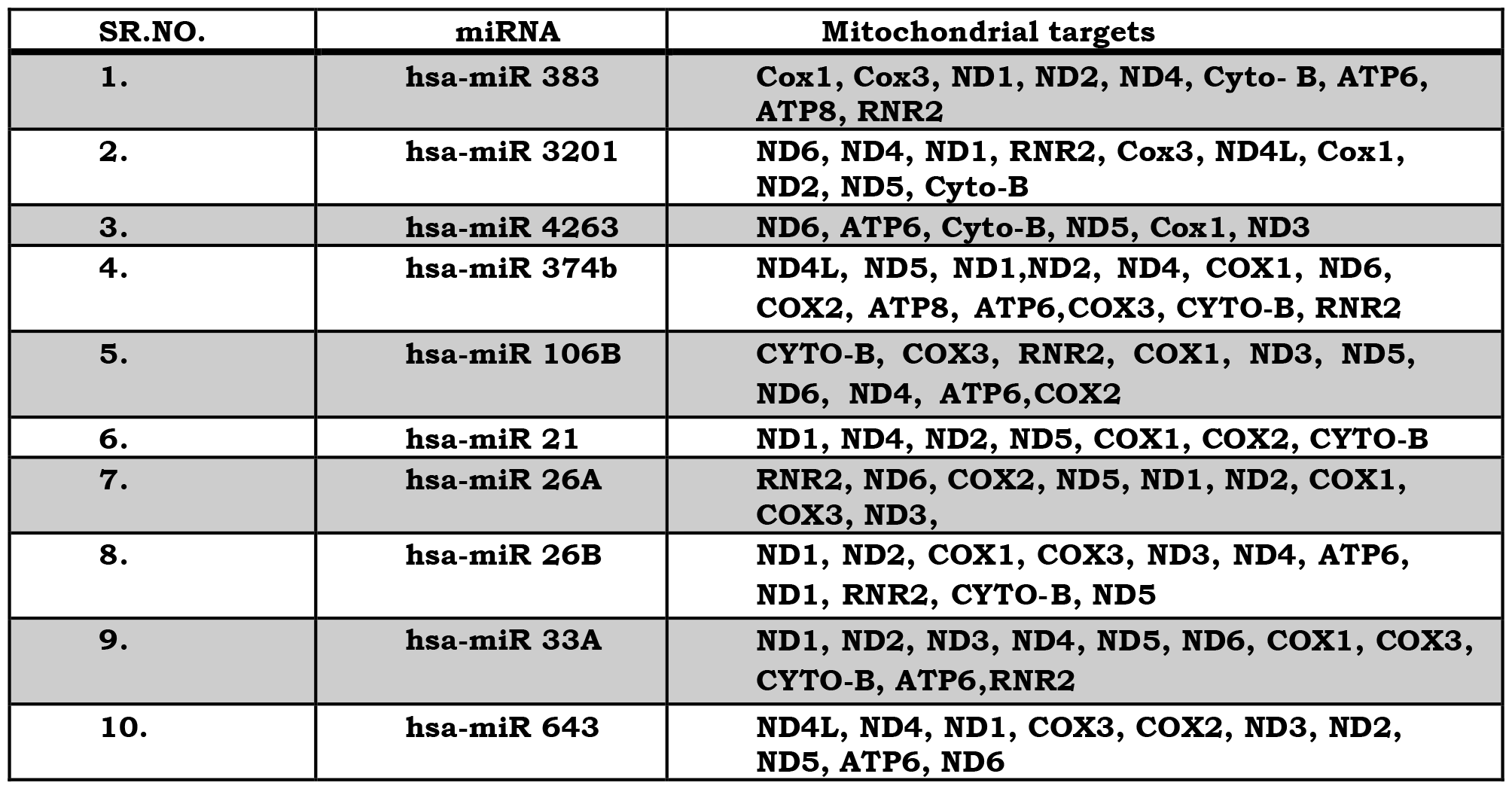
Candidate miRNAs and their predicted mitochondrial target genes. The target prediction analysis was done using miRDB, MiRanda, Target scan, miRbase and microRNA.org.

Following target prediction study, we performed scoring analysis to select the highest scored mitochondrial gene targeted by individual microRNA. To achieve this, the predicted targets of each of the 5 bioinformatic database (i.e. mirBase, miRanda, Target scan, microRNA.org, and miRDB), were compared for the presence & absence of the specific target, in each of these algorithms. And the highest scored mitochondrial target genes were selected for each of the candidate micro RNA, for the target validation study.

**(Table 3.3)**

**Table 3.3:** Scoring of mitochondrial genes targeted by candidate microKNA. 1 He targets predicted by each individual algorithm was compared for presence or absence of specific target gene and was scored based on the number of data bases which predicted the gene as target for the miRNAs. The highest scored gene was selected for target validation study. A, B, C, I), E, E, G, 11 and I represents the scoring chart of miR 4263, 383, 21, 26b, 33a, 643, 3201, 374b, 106b and 26a respectively.

| miRNA | Mt genes (Targets) | miRDB | miRanda | miRbase | MicroRNA.org | Target scan | TOTAL |
| --- | --- | --- | --- | --- | --- | --- | --- |
| hsa-miR-4263 | Cox1 |  |  |  | * |  | 1 |
|  | Cyto. B |  |  | * |  |  | 1 |
|  | ND3 |  |  | * |  |  | 1 |
|  | ND4L |  |  | * |  |  | 1 |
|  | **ND6** | * | * | * | * | * | 5 |
|  | ATP6 |  |  |  | * |  | 1 |
|  | RNR2 |  |  | * | * |  | 2 |

A.
| miRNA | Mt genes (Targets) | miRDB | miRanda | miRbase | MicroRNA.org | Target scan | TOTAL |
| --- | --- | --- | --- | --- | --- | --- | --- |
| hsa-miR-383 | **Cox1** | * | * | * | * | * | 5 |
|  | Cox3 |  |  | * |  |  | 1 |
|  | Cyto. B |  |  |  | * |  | 1 |
|  | **ND4L** |  | * | * | * | * | 4 |
|  | ND2 | * |  |  |  |  | 1 |
|  | ND6 |  |  | * | * | * | 3 |
|  | **ATP6** | * |  | * | * | * | 4 |
|  | ATP8 |  |  |  | * |  | 1 |
|  | RNR2 | * |  |  |  |  | 1 |

B.
| miRNA | Mt genes (Targets) | miRDB | miRanda | miRbase | MicroRNA.org | Target scan | TOTAL |
| --- | --- | --- | --- | --- | --- | --- | --- |
| hsa-miR-21 | **Cox1** | * | * | * | * | * | 5 |
|  | Cyto-B |  |  |  | * |  | 1 |
|  | ND4 |  |  |  | * |  | 1 |
|  | ATP6 |  |  | * |  |  | 1 |
|  | ND6 |  |  |  | * |  | 1 |

C.
| miRNA | Mt genes (Targets) | miRDB | miRanda | miRbase | MicroRNA.org | Target scan | TOTAL |
| --- | --- | --- | --- | --- | --- | --- | --- |
| hsa-miR-26b | Cox1 |  |  |  | * |  | 1 |
|  | Cox3 |  |  |  | * |  | 1 |
|  | Cyto. B |  |  |  | * |  | 1 |
|  | ND1 |  |  |  | * |  | 1 |
|  | ND2 |  |  | * |  |  | 1 |
|  | ND3 |  |  |  | * |  | 1 |
|  | ND4 | * |  |  |  |  | 1 |
|  | ND5 |  |  |  | * |  | 1 |
|  | ATP6 |  |  |  | * |  | 1 |
|  | **RNR2** | * | * | * | * | * | 5 |

D.
| miRNA | Mt genes (Targets) | miRDB | miRanda | miRbase | MicroRNA.org | Target scan | TOTAL |
| --- | --- | --- | --- | --- | --- | --- | --- |
| hsa-miR-33a | Cox1 |  |  | * | * |  | 2 |
|  | Cox3 |  |  |  | * |  | 1 |
|  | Cyto. B | * |  | * | * |  | 3 |
|  | ND1 |  | * | * | * |  | 3 |
|  | ND2 |  |  | * |  |  | 1 |
|  | ND3 |  |  |  | * |  | 1 |
|  | **ND4** |  | * | * | * | * | 4 |
|  | **ND5** | * | * | * | * | * | 5 |
|  | ND6 |  |  | * |  |  | 1 |
|  | ATP6 |  |  |  |  | * | 1 |
|  | RNR2 |  | * | * | * | * | 4 |

E.
| miRNA | Mt genes (Targets) | miRDB | miRanda | miRbase | MicroRNA.org | Target scan | TOTAL |
| --- | --- | --- | --- | --- | --- | --- | --- |
| hsa-miR-643 | Cox2 |  |  | * | * | * | 3 |
|  | Cox3 |  |  | * |  |  | 1 |
|  | ND1 |  |  |  | * |  | 1 |
|  | ND2 | * |  | * |  |  | 2 |
|  | ND3 |  |  |  | * |  | 1 |
|  | ND4 |  |  | * |  |  | 1 |
|  | ND4L |  |  | * |  |  | 1 |
|  | ND5 |  |  | * | * |  | 2 |
|  | ND6 | * |  | * |  |  | 2 |
|  | ATP6 |  |  | * | * |  | 2 |

F.
| miRNA | Mt genes (Targets) | miRDB | miRand a | miRbase | MicroRNA. org | Target scan | TOTAL |
| --- | --- | --- | --- | --- | --- | --- | --- |
| hsa-miR-3201 | Cox1 |  | * | * | * |  | 3 |
|  | Cox3 |  |  |  | * |  | 1 |
|  | Cyto. B |  |  |  | * | * | 2 |
|  | ND1 |  |  | * |  |  | 1 |
|  | ND2 |  |  | * | * |  | 2 |
|  | ND4 |  |  |  |  |  | 1 |
|  | **ND4L** |  | * | * | * | * | **4** |
|  | ND5 |  |  | * | * |  | 2 |
|  | ND6 |  |  |  |  | * | 1 |
|  | RNR2 |  |  |  | * |  | 1 |

**F.**
| miRNA | Mt genes (Targets) | miRDB | miRanda | miRbase | MicroRNA. org | Target scan | TOTAL |
| --- | --- | --- | --- | --- | --- | --- | --- |
| hsa-miR-374B | Cox1 | * |  |  | * |  | 2 |
|  | Cox2 |  |  | * |  |  | 1 |
|  | Cox3 |  |  | * |  |  | 1 |
|  | Cyto. B |  |  | * |  |  | 1 |
|  | ND1 |  | * |  |  | * | 2 |
|  | ND2 |  |  | * | * |  | 2 |
|  | ND4 |  |  | * |  | * | 2 |
|  | **ND4L** |  | * | * | * | * | **4** |
|  | ND5 | * |  | * | * |  | 3 |
|  | ND6 |  |  | * |  |  | 1 |
|  | ATP6 |  |  |  | * |  | 1 |
|  | ATP8 |  |  |  |  | * | 1 |
|  | RNR2 | * |  | * | * |  | 3 |

**G.**
| miRNA | Mt genes (Targets) | miRDB | miRanda | miRbase | MicroRNA. org | Target scan | TOTAL |
| --- | --- | --- | --- | --- | --- | --- | --- |
| hsa-miR-106B | Cox1 |  |  | * | * |  | 2 |
|  | Cox2 |  |  | * |  |  | 1 |
|  | Cox3 |  |  | * | * | * | 3 |
|  | **Cyto. B** | * |  | * | * | * | **4** |
|  | ND3 | * |  |  | * |  | 2 |
|  | ND4L |  |  |  | * |  | 1 |
|  | ND5 |  |  | * | * |  | 2 |
|  | ND6 | * |  |  |  |  | 1 |
|  | ATP6 | * |  |  |  |  | 1 |
|  | RNR2 |  |  | * | * |  | 2 |

**H.**
| miRNA | Mt genes (Targets) | miRDB | miRanda | miRbase | MicroRNA. org | Target scan | TOTAL |
| --- | --- | --- | --- | --- | --- | --- | --- |
| hsa-miR-26a | Cox1 |  |  |  | * |  | 1 |
|  | Cox2 |  |  |  | * |  | 1 |
|  | Cox3 |  |  |  | * |  | 1 |
|  | ND1 | * |  |  |  |  | 1 |
|  | ND2 |  |  | * | * |  | 2 |
|  | ND3 |  |  |  | * |  | 1 |
|  | ND5 | * |  | * |  |  | 2 |
|  | ND6 | * |  |  |  |  | 1 |
|  | RNR2 |  |  | * | * |  | 2 |

### Comparative analysis of expression pattern of the candidate miRNAs, in clinical samples of HCC

The bioinformatic study of the differential expression pattern of candidate miRNAs in the clinical samples of hepatocellular carcinoma (HCC) was performed by using Gene expression Omnibus database (Accession ID-GSE123972). To achieve this 47 clinical samples of HCC, at initial and advance stage, were compared to check the expression pattern of the candidate microRNAs. The study revealed that miRNA 383, 21 and 106b were differentially expressed in HCC patients when compared from among the 10 candidate miRNAs selected. A total of 1119 microRNAs were analyzed, out of which 546 microRNAs were upregulated while 217 were significantly downregulated (Tan C. et. al, 2019, Accession ID-GSE123972).

**(Figure 3.1)**

**Figure 3.1:**
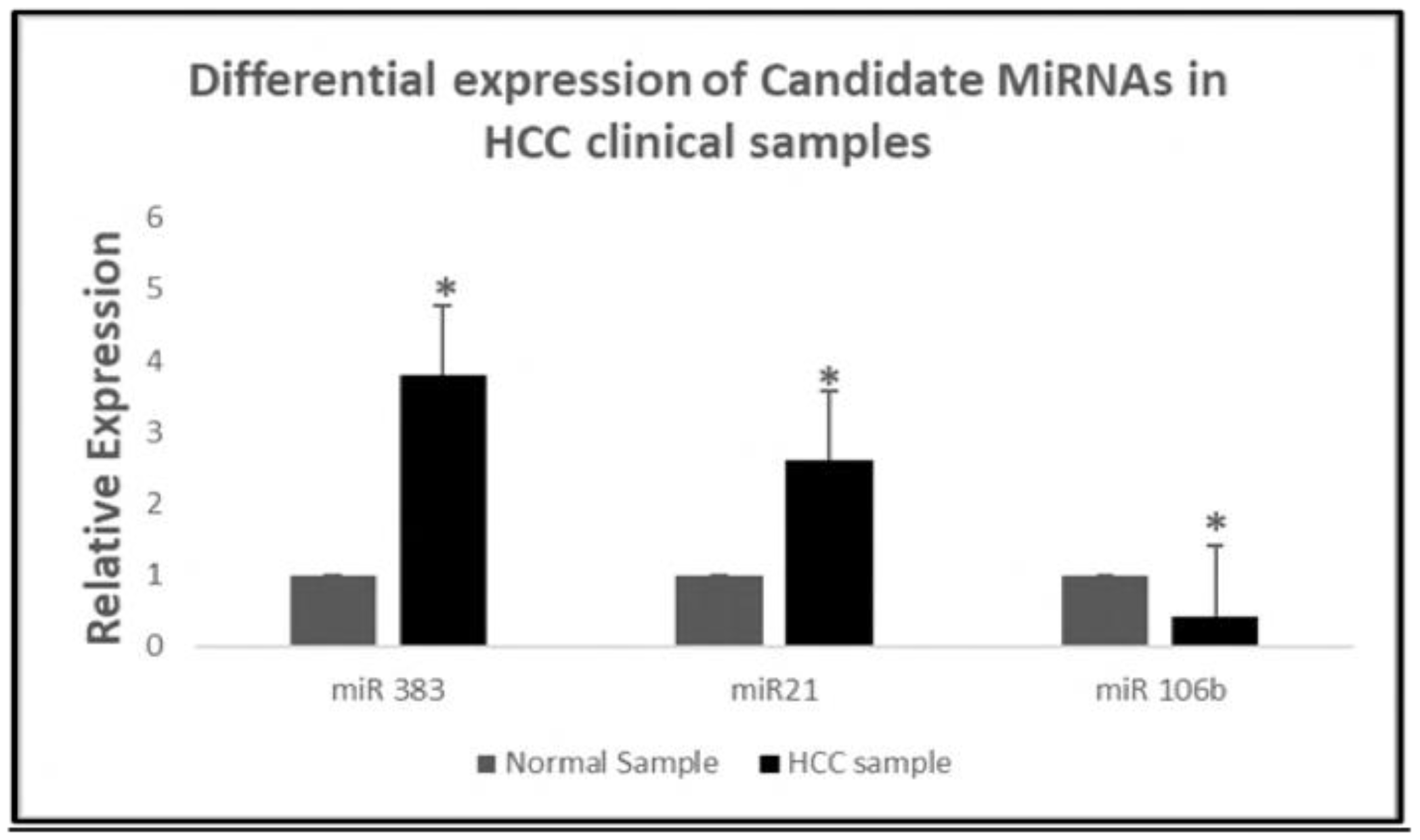
Relative expression of microRNA 383, 21 and 106b in clinical sample of HCC. 1119 micro RNAs from plasma samples of HCC, at initial and advance stage, were compared to check the expression pattern of candidate microRNAs. Results suggested that miR 21, 106b and 383 were differentially expressed in HCC. Results presented are average of three experiments ± SEM each done at least in triplicate. p<0.05. *Statistically significant when compared to control.

### MicroRNAs enrichment in HepG2 and WRL cells

The result of qRT PCR revealed that all the 10 candidate microRNAs were differentially expressed in the HepG2 cells when compared with WRL cells. 3 out of 10 microRNAs i.e. (miR 21, 383, 3201) were significantly down regulated; rest all microRNAs showed a higher expression level.

**(Figure 3.2)**

**Figure 3.2:**
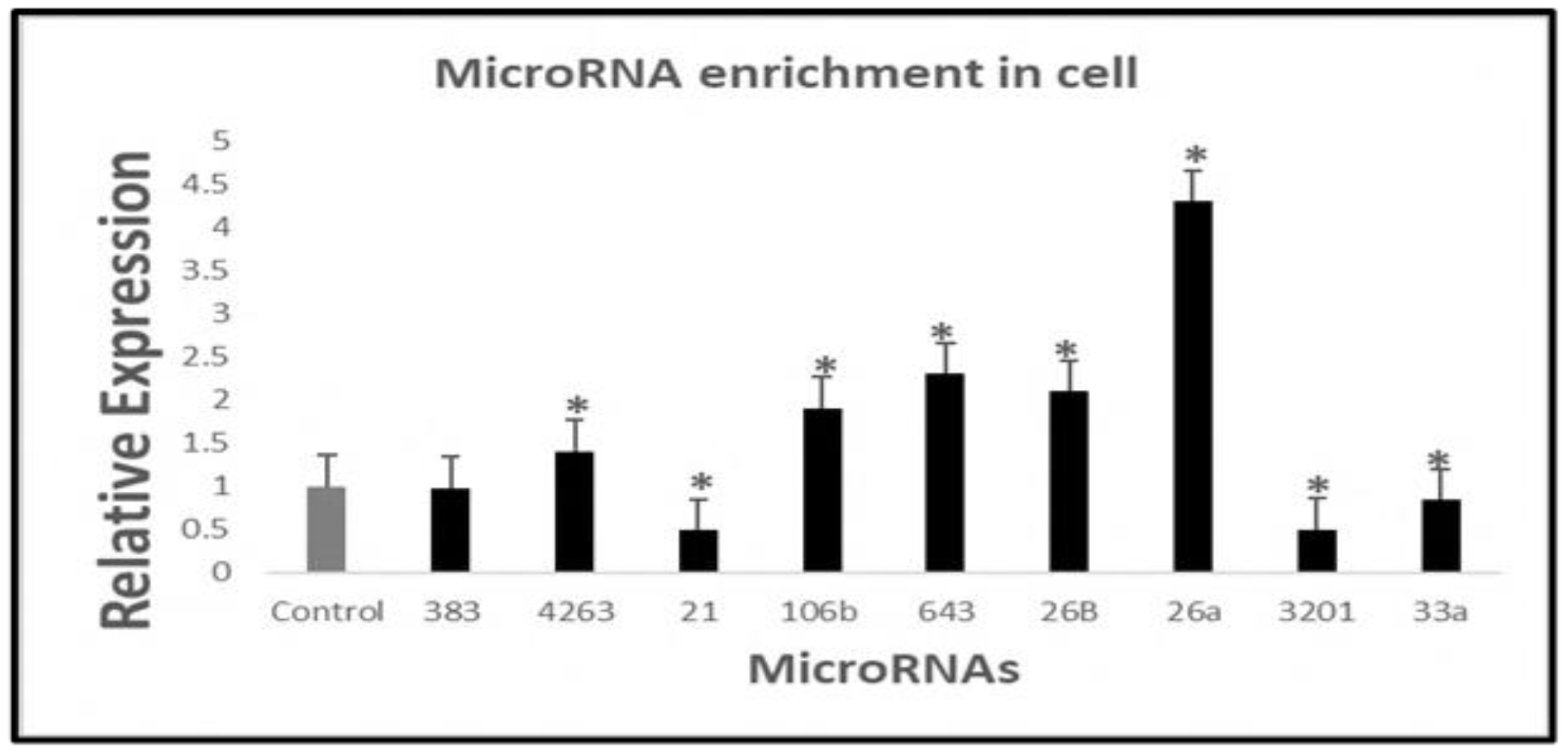
Relative expression pattern of the candidate microRNAs. RT-PCR analysis was done in HepG2 keeping WRL cells as control, to check the expression pattern of the candidate microRNAs. Ratio of expression was calculated using the expression level in WRL as 1. The results revealed that the expression of miR 21, 383. and 3201 was low in HepG2. Results presented are average of three experiments ± SEM each done at least in triplicate, p<0.05. * Statistically significant when compared to control.

### Differential expression pattern of the miRNAs in mitochondria

Following the analysis of miRNA expression in cell, we checked the expression level of these microRNAs in the mitochondria of HepG2 and WRL cells. For, this, mitochondria was isolated from WRL and HepG2 cell and the purity was checked by the western blot analysis using mitochondria specific marker. Further, nuclear and cytoplasmic contamination was ruled out by checking nucleus and cytoplasm specific marker in the mitochondrial pellet. The results revealed that there is a differential pattern of expression of the 10 candidate microRNA in mitochondria isolated from HepG2 cell when compared with WRL. 8 out of the 10 micro RNAs were found to be significantly lower in the mitochondria and 2 were significantly high.

**(Figure 3.3 and figure 3.4)**

**Figure 3.3:**
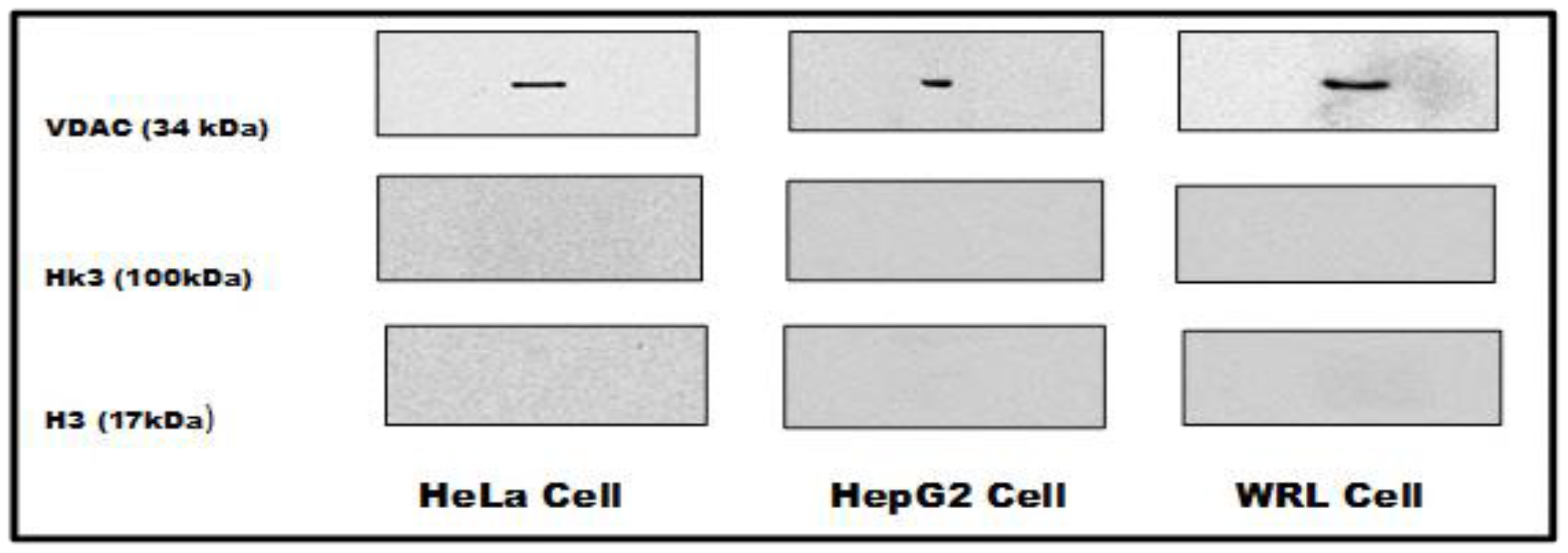
Mitochondrial pellet was free of cytoplasmic and nuclear fractions. Western blot analysis was performed to check the purity of the mitochondrial pellet by checking with mitochondrial marker VDAC, cytoplasmic marker HK3 and nuclear marker protein H3. The results revealed that mitochondrial pellet was free from cytoplasmic and nuclear fractions, as only VDAC gave the protein band, whereas HK3 and H3 did not give any visible b and. Results presented are average of three experiments ± SEM each done at least in triplicate, p<0.05. * Statistically significant when compared to control.

**Figure 3.4:**
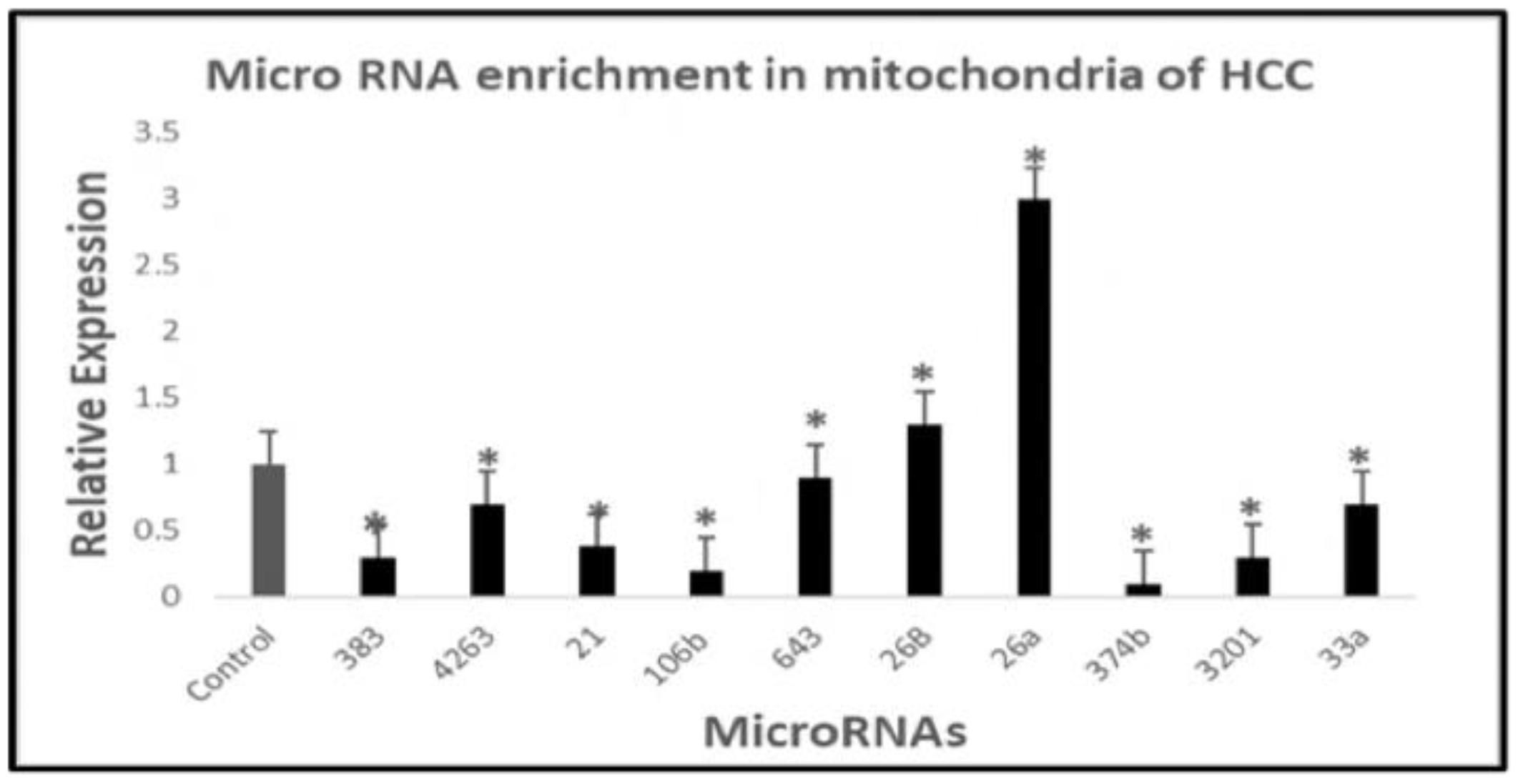
Expression pattern of the candidate microRNAs in mitochondria. RT-PCR analysis was performed to check the expression pattern of the candidate microRNAs in the mitochondria of HepG2 cells keeping WRL cells as control. Ratio of expression was calculated using the expression level in WRL as 1. The results revealed that the expression of miR 26a and 26b was high in the mitochondria of HepG2, whereas the expression of rest of the candidate microRNAs were found low in the mitochondria of HepG2 cells. Results presented are average of three experiments ± SEM each done at least in triplicate, p<0.05. * Statistically significant when compared to control.

#### Expression pattern of mitochondrial genes were high in the mitochondria

Following analysis of differential expression of candidate microRNAs in the mitochondria, we checked the expression levels of their target mitochondrial genes and found that levels of all the 5 candidate mitochondrial target genes were high in the mitochondria of HepG2 cells. The results shown a 170 fold higher levels Cyto-B in the mitochondria of HepG2 cells, the levels of Cox1, ATP6, NDL-4 and ND6 was found 16, 8, 42 and 17 fold higher respectively.

**(Figure 3.5)**

**Figure 3.5:**
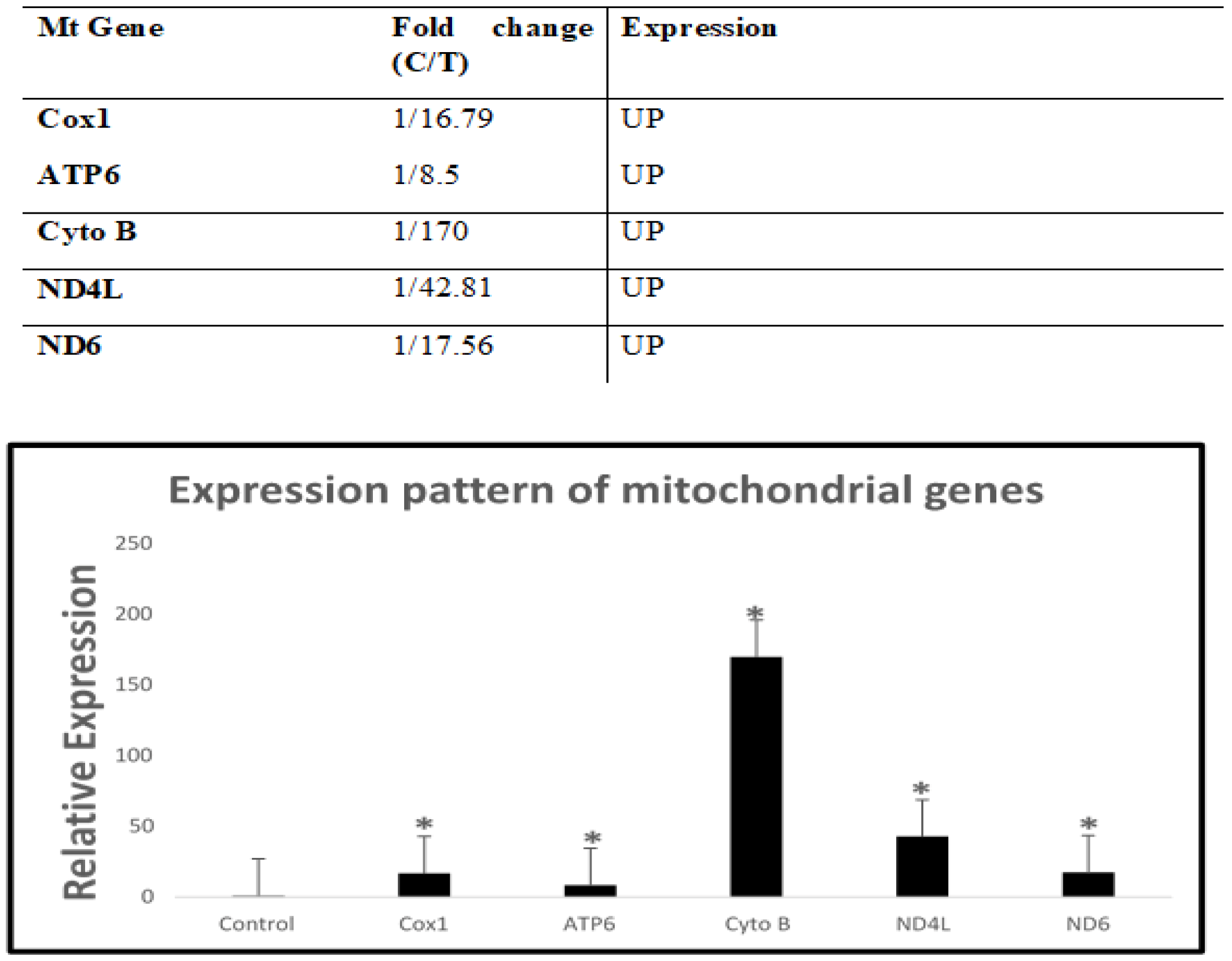
Relative expression pattern of the mitochondrial genes. The mitochondria was isolated from HepG2 & WRL cells and the RT-PCR analysis was performed to check the expression pattern of mitochondrial genes. The results suggested that all the 5 target mitochondrial genes were at high levels in the mitochondria of HepG2 cells when compared with WRL cells. A.) Ratio of expression of mitochondrial genes in HepCr2 and WRL cells (calculated using the expression level in WRL as 1. B.) Relative expression of the mitochondrial genes. Results presented are average of three experiments ± SEM each done at least in triplicate, p<0.05. * Statistically significant when compared to control.

## 4. Discussion

Mitochondria is often denoted as the power house of the cells as they synthesis ATP, required for normal cellular functioning and this makes them necessary for eukaryotic life (Mcbride et.al, 2014). Mitochondria also takes part in breaking down sugars and lengthy chains of fatty acids, synthesizing steroids and lipids, and performing several other vital processes for human life (Amaloha C et. al 2023). The stability and proper functioning of the mitochondria is crucial for the cells survivability and any alteration in the mitochondrial machinery could lead to several diseased conditions (Khan T. et. al, 2022, Li A et.al, 2020). The malfunctioning of the mitochondria was first reported by Warburg, 70 years ago, where he noticed that tumors produces an excess of lactate in the presence of oxygen, naming it aerobic glycolysis or Warburg effect (Pascale et.al, 2020, Liberty M et.al, 2016).

Mutations in the mtDNA or suppression of mitochondrial genes have been reported to be the main cause of mitochondrial dysfunction in a variety of cancers (Wang SF et.al, 2023). These changes appear to modify the mitochondrial metabolism, start tumor progression, and enable malignant cells to adapt to their changing environment (Elkholi et.al, 2014). Mitochondrial function is also impacted by the activation of oncogenes and various signaling pathways like PI3K–PTEN–AKT pathway (Lien EC et. al, 2016), which causes the mitochondrial metabolism to change from oxidative to glycolytic (Liu C et al., 2021, Yan Hi et.al, 2021). Despite having their own genome, the vast majority of mitochondrial proteins are imported into the mitochondria by the nuclear genome (Pfanner N et.al, 2021). Therefore, changes in the expression of mitochondrial genes as well as nuclear genes, necessary for mitochondrial function may result in the modulation of mitochondrial metabolism (Zhou Q et. al, 2021, Douglas C, 2010).

MicroRNA is a class of non-coding RNA which possess the ability to regulate the expression pattern of a variety of genes (He L et.al, 2004), playing role in various vital processes involved in normal cellular functioning (O Brien et.al, 2018). The microRNAs which are either coded by the mitochondrial genome or by nuclear genome and imported to the mitochondria, regulating the expression pattern of the mitochondrial genes are called MitomiRs (Purohit P et. al, 2020). Various MitomiRs have been reported to alter the mitochondrial metabolism leading to diseases like neurodegenerative, cardiac, cancer etc. (John A et.al, 2020, Duarte F et. al, 2015).

For this study, we selected 10 most important microRNAs, reported to be involved in the various types of cancer. The selection of the candidate microRNA was done, based on their nature (oncogenic or tumor suppressive), their involvement in common cancer and which have not been yet reported to be involved in regulation of electron transport chain. To achieve this, we used Pubmed database and performed a detailed survey of all the studies done till date involving the candidate microRNAs with our search criteria and all together 215 publications were retrieved and reviewed. With the literature review, we found that the candidate microRNAs were seen in regulating various genes involved in the process of cancer initiation and progression; however we could not find any report on their role in modulating the mitochondrial machinery particularly, electron transport chain.

Next we checked, if these microRNAs have targets on the crucial mitochondrial genes, by target prediction analysis using 5 different bioinformatic algorithms i.e. miRDB, microRNA.org, Target scan, miRBase, miRANDA. The results revealed that the candidate microRNAs have targets on several important mitochondrial genes involved in the ETC. Following this, we performed scoring analysis, where we compared the predicted targets of each algorithm, for the presence or absence of specific mitochondrial target gene and the highest scored genes were selected for target validation study.

Parallely, we performed a comparative study of the microRNAs involved in HCC using 47 clinical samples of HCC with gene expression omnibus database of NCBI. The result shown that, 3 microRNAs (miR 383, miR21, miR 4263) out of 10 candidate microRNAs were differentially expressed in the clinical samples of HCC patients when compared with the healthy individual.

Various studies suggest that microRNAs are differentially expressed in cancer. So, we performed qRT PCR analysis to check the relative expression pattern of the candidate microRNAs in HepG2 by comparing it with WRL cells. The results of this study, showed that levels of miR 21, 33a, 3201, and 383 were significantly low, while miR 26a, 26b, 4263, 643 and 106b were found high in HepG2 cells. Following this, we checked the microRNA enrichment in the mitochondria of HepG2 cells by RT-PCR. To achieve this, mitochondria was isolated and purity was checked with western blot analysis using mitochondrial marker protein VDAC, followed by RT-PCR. The results indicated that the levels of microRNA 383, 4263, 21, 106b, 643, 374b, 3201 and 33a were found to be significantly low in the mitochondria of HepG2, while the level of miR 26a and 26b was high. The target prediction analysis revealed that these down regulated microRNAs possesses targets on the important mitochondrial genes, so next we checked the impact of their down regulation on the expression of mitochondrial genes by qRT-PCR. The results revealed that the expression pattern of the mitochondrial genes was significantly high in the HepG2 cells when compared with WRL cells.

This inverse relation of the expression of microRNAs to their mitochondrial target genes, made us to hypothesize that since the expression of the microRNA which have targets on these mitochondrial genes is low in the mitochondria, so the mitochondrial target genes show a higher expression level and if we would over express the candidate micro RNAs, the expression level of their target genes will go down, and also, this down regulation of the mitochondrial genes could alter the mitochondrial machinery and may lead to carcinogenesis.

In contrary to the reported levels of miR 383 & miR 21 as over expressed in most of hepatocellular carcinoma, we found a model of HepG2, where the levels of miRNA was either equal or lower than the non-cancerous cell line (WRL). This provided us an apt model system to check the effect of these microRNAs (which are over expressed in tumor condition) on mitochondrial gene targeting & mitochondrial function when over expressed.

## Acknowledgment

We acknowledge Indian Council of Medical Research, Ministry of Health, Govt. of India for the financial assistance in form SRF and Kerala state council for Science Technology & Environment, Govt. of Kerala for fellowship in the form of JRF and SRF to Mr. Ashutosh K. Maurya. We also acknowledge Central University of Kerala for providing all the necessary facilities to carry out this research work.

## Author Contributions

The authors confirm contribution to the paper as follows: Study conception and design: VBSK, Bioinformatics and wet lab work: AKM. Analysis of clinical data: RP. All authors reviewed the results and approved the final version of the manuscript.

## Conflicts of Interest

The authors declare that they have no conflicts of interest to report regarding the present study.

## SUPPLEMENTAL DATA

## Notes

### Competing Interest Statement

The authors have declared no competing interest.

